# Mapping the interactome of human tRNA methyltransferase TRMT1 using dual proximity labeling

**DOI:** 10.64898/2026.05.18.725941

**Authors:** Angel D’Oliviera, Sophie Olson, Harrison Bernhard, Yanbao Yu, Jeffrey S. Mugridge

**Affiliations:** University of Delaware, Department of Chemistry & Biochemistry, Newark DE 19716

## Abstract

Transfer RNA methyltransferase 1 (TRMT1) installs N2-methylguanosine and N2,N2-dimethylguanosine modifications at position 26 of mammalian tRNAs, supporting tRNA structure, translation, and cellular response to redox stress. However, the local environment and interactome of TRMT1 in the cell is poorly defined. Here, we use APEX2-based proximity labeling of the N- and C-terminus of TRMT1, coupled with label-free quantitative proteomics to map candidate TRMT1-proximal proteins in HEK293T cells. Mass spectrometry data was acquired using both data-independent acquisition (DIA) and data-dependent acquisition (DDA) methods, and it was found that DIA substantially increased proximity proteome coverage, reproducibility, and the number of significantly enriched candidate hits compared to the DDA method. N- and C-terminal APEX2-TRMT1 constructs captured largely overlapping proteomes, suggesting the dual-labeling strategy provides a robust map of proximal proteins. Analysis of the significant TRMT1-proximal proteins reveals enrichment in RNA processing and ribonucleoprotein-associated factors, in addition to hits connected to tRNA modification, tRNA biogenesis, and redox-associated biology. These data provide a proteome-scale view of TRMT1-associated cellular proteins and environments, and lay the groundwork for future validation of functional TRMT1 interaction networks.

**Significance:**

- Fusing APEX2 enzyme to both N-terminal and C-terminal of the bait enhanced the sensitivity for identification of protein interactions.
- Combining APEX2-based endogenous labeling with DIA mass spectrometry increases reproducibility and depth of proximity proteome.
- The study provides a rich source of potential interacting or proximally close proteins to TRMT1, which warrants further validation studies.

## Introduction

The cell uses diverse chemical modifications on transfer RNA (tRNA) molecules to help regulate tRNA function across all domains of life [1–3]. Of the over 150 different RNA modifications discovered in all types of organisms, about 80% have been identified on tRNA and each tRNA molecule in eukaryotes has an average of 13-14 modifications that can directly impact their structure, stability, aminoacylation, codon-anticodon base pairing, and translation fidelity [1–9]. Additionally, multiple modifications can work in tandem to generate cumulative effects on tRNA function and physiology [2,7]. Mutations in tRNA modifying enzymes or dysregulation of tRNA modification pathways often lead to significant defects in protein translation and protein homeostasis, and are closely linked to the development and progression of human diseases including neurodegenerative disorders, cancers, and mitochondrial diseases [3–6].

The most common type of tRNA modification is nucleobase methylation, which can affect tRNA structure and stability, protein-tRNA interactions, translation regulation, metabolic processing in response to cellular condition/stress, and other cellular processes imperative for maintaining cellular homeostasis [10,11]. The vast majority of methyl modifications on tRNA are installed by *S*-adenosyl-*L*-methionine (SAM)-dependent methyltransferases, which use cofactor SAM as a methyl donor [12–15]. The diverse SAM-dependent methyltransferase superfamily is classified into mainly five distinct structural families or classes (I-V), determined primarily by enzyme fold [16–18]. The majority of tRNA methyltransferases are members of class I (characterized by a Rossman-fold domain) and IV (characterized by a trefoil knot feature), and typically mono-methylate nucleobase nitrogen atoms at endocyclic (e.g. N1, N3, N7) positions or mono-/di-methylate nitrogen or oxygen atoms at exocyclic (e.g. guanosine N2, adenosine N6, ribose 2′-O) positions [11,12]. Examples of the most abundant and conserved methyl modifications on tRNA at commonly modified atoms include m5C, m1A, m3C, m6A, and m2G, which play key roles in stability, translational efficiency, global protein synthesis, and stress responses [10,12,19].

tRNA methyltransferase 1 (TRMT1) catalyzes installation of the N2,N2-dimethylguanosine (m2,2G) or N2-methylguanosine (m2G) modification on the guanosine base at position 26 in mammalian tRNAs [20]. m2,2G modifications play significant roles in regulation of tRNA conformation and function by providing structural stability to the tRNA region where the D-arm and anticodon stem meet, preventing the misfolding of tRNA by inhibiting undesirable base pair interactions [1–9,21]. Loss of m2,2G26 on tRNA results in impaired global translation, slowed cellular growth, defects in oxidative stress response, and altered tRNA processing in mitochondria [7,22]. The phenotypes of m2G loss are less evident, but likely include mild impacts on translation and tRNA processing [7]. Cells which lack or have pathogenic variants of TRMT1 are more susceptible to oxidative stress, particularly in neurons, and human TRMT1 variants are associated with intellectual disabilities and various neurological disorders including epilepsy and congenital microcephaly [21,22].

Most cytosolic tRNA molecules with G at position 26 have the m2,2G modification, while very few possess m2G (tRNA-iMet and tRNA-Val are two known examples), and others have unmodified G26 [7,21,23]. In mitochondria, this trend is seemingly reversed, with most G26 containing tRNAs having the m2G modification and only one, tRNA-Ile, having the m2,2G modification. This suggests that TRMT1-mediated tRNA methylation is selective and regulated within the cell, which may in part be controlled by protein-protein interactions. The activity and targeting of many other methyltransferases are regulated by protein interactions, for example tRNA m^7^G methyltransferase METTL1 forms a complex with WDR4 to enhance substrate binding, and tRNA m^3^C methyltransferase METTL6 forms a complex with seryl-tRNA synthetase to impart substrate selectivity [24–26]. Therefore, mapping the TRMT1 protein interactome is a key step toward understanding the regulation of TRMT1 methylation activity and selectivity in the cell. Proximity labeling coupled with mass spectrometry-based quantitative proteomics has been a powerful tool to investigate and identify interacting proteins in native cellular environments [27]. However, targeted, proteome-wide investigation focused on TRMT1 interactors has not previously been carried out. Here, for the first time, we employed proximity labeling based quantitative proteomics to map the spatial and protein interaction networks of TRMT1 in living cells. We constructed APEX2-fusions to either the N-terminus or C-terminus of TRMT1, and utilized this dual-labeling approach to enhance the sensitivity and accuracy of the mapped interactions. We also employed both data-independent acquisition (DIA) and data-dependent acquisition (DDA) mass spectrometry, and carried out head-to-head comparisons of the two pulldown complexes to assess their performance for proximity proteomics analysis. The comprehensive dataset obtained from this study serves as a valuable resource for future studies of TRMT1 and tRNA modification biology.

## Materials and Methods

### Construct Design and Cloning

To generate TRMT1-APEX2 fusions, the full-length human gene sequence TRMT1 (NM_001136035.4) and a codon-optimized 3x GGGS linker (GGAGGAGGAGGAAGC nucleotide sequences repeated three times consecutively) were inserted into the APEX2-pLX304 plasmid using Gibson Assembly, on either terminus of APEX2 to generate a TRMT1-3xGGGS-APEX2 and APEX2-3xGGGS-TRMT1 construct. The lentiviral packaging plasmid pCMV-dR8.2 dvpr (Addgene #8455), envelope protein plasmid for producing lentiviral particles pCMV-VSV-G (Addgene #8454), and APEX2-pLX304 plasmid (Addgene #92158) plasmids were sourced from Addgene.

All final plasmids were transformed into NEB 5-alpha Competent *E. coli* (ref: C2987). To obtain large quantities of DNA for mammalian transfection, a single colony for each construct was grown overnight in 500 mL LB media with 50 µg/mL ampicillin, at 37 °C with shaking at 200 rpm. DNA was prepared using Qiagen Plasmid Maxi Prep kit (ref: 12162) following the commercial protocol to obtain total DNA yields greater than 500 µg for future HEK293T transfections.

### HEK293T Adherent Cell Culture

Human embryonic kidney (HEK) 293T cells were obtained from ATCC (ref: CRL-3216). All lines were cultured in growth medium (Dulbecco’s Modified Eagle’s Medium (DMEM) supplemented with 10% fetal bovine serum (FBS) and penicillin (100 U/mL) + streptomycin (100 µg/mL)), at 37 °C under 5% CO_2_ in tissue culture treated 100 mm standard mammalian cell culture dishes. Every 2-3 days the growth media was replaced, and the cells were subsequently expanded after reaching 80-90% confluency (approximately 5-7 days).

To passage, the growth media was removed, and cells were washed with prewarmed PBS. After removing the PBS, 0.25% Trypsin in HBSS with 0.2 g/L EDTA was mixed with PBS at a 1:2.5 ratio and incubated with washed cells at 37°C for approximately 1-2 min to detach adherent cells, after which, a 3-fold dilution with growth media was added for trypsin inactivation. The cells were mixed in a homogenous suspension and diluted 1:20 with growth media and added into a new cell culture dish.

### Generation of APEX2-fusion HEK293T Stable Cell Lines

Low passage HEK293T cells were cultured to 90% confluency and the media was replaced with fresh DMEM containing 10% FBS on the day of transfection. Lentiviral vectors were produced using a Lipofectamine 3000 transfection method (ref: L3000015). For each sample, 30 µL of Lipofectamine 3000 reagent was diluted in 3 mL Opti-MEM I Reduced Serum Media. For each transfection, the individual APEX2 construct (6.3 µg) was combined with the lentiviral packaging (pCMV-dR8.2 dvpr, 5.7 µg) and envelope (pCMV-VSV-G, 600 ng) plasmids, and subsequently mixed with 30 μL P3000 Reagent and 3 mL of Opti-MEM I Reduced Serum Media. This solution was mixed 1:1 with Lipofectamine 3000 reagent solution and incubated for 30 minutes at 25 °C. The plasmid and Lipofectamine complexes were then incubated with HEK293T cells at 37 °C, 5% CO_2_. After 3 hours, the transfection mixture was replaced with growth medium and the cells were incubated for 48 hours, after which, the supernatant containing the lentivirus was harvested and filtered using a 0.45 µm filter. This lentiviral-containing media was then used for transduction at a 1:200 dilution into fresh HEK293T cells at approximately 30% confluency. Following 72 hours of incubation, growth media containing 8 µg/mL Blasticidin S HCl was replaced daily and cells were passaged every 2-3 days. 14 days post-transduction, stable expression of APEX2-fusion constructs was verified through western blot. Stocks of the stable cell lines in cryovials were made with 1 × 10^6^ cells/mL in complete DMEM media containing 5% DMSO, slow frozen at -80 °C, overnight and stored in liquid nitrogen. Stable cell lines were generated for the N-terminal APEX2-TRMT1 fusion, the C-terminal TRMT1-APEX2 fusion, and the APEX-only construct.

### Western Blot

To verify the expression of our APEX2 fusion stable constructs, HEK293T cells were trypsinized as described above, and the resulting cell suspension was spun down at 125 x g for 3 minutes. Cell pellets were washed in ice-cold PBS 3 times and collected at 125 x g for 3 minutes, aspirating PBS in between each wash, following which the pellets were flash frozen using liquid nitrogen and stored at -80 °C. Cell pellets were thawed on ice and lysed with 500 µL of Lysis Buffer (150 mM NaCl, 1% Triton X-100, 0.5% Sodium Deoxycholate, 0.1% SDS, 50 mM Tris HCl pH 8.0, 10 mM ascorbate, 5 mM Trolox, cOmplete Mini EDTA-free Protease Inhibitor Cocktail (ref: 11836170001). Lysate was spun at 12,000 x g, and the supernatant was collected and boiled in SDS-PAGE sample buffer and loaded (10 µL) onto a Bio-Rad TGX 4-15% polyacrylamide gel and run for 30 minutes at 180 V. Gels were blotted onto PVDF membranes using a Bio-Rad Trans-blot Turbo for 7 minutes at 2.5 Amps. The blot was incubated in blocking solution (5% non-fat milk in 1X Tris-Buffered Saline with 0.1% Tween) at room temperature for 1 hour. All antibodies were diluted in blocking solution. TRMT1-specific antibodies corresponding to amino acid regions 460-659 (Invitrogen Rabbit Anti-TRMT1 (ref: PA5-96585)) and 609-659 (Bethyl Laboratories Rabbit Anti-TRMT1 (ref: A304-205A)) and Anti-APEX2 antibodies (Abcam ref: ab222414) were each utilized for primary antibody staining at 1:2,000 and were stained overnight at 4°C. The blot was rinsed three times with 1X Tris-buffered Saline with 0.1% Tween after primary antibody staining. Invitrogen Goat anti-Rabbit IgG (H+L) Secondary Antibody conjugated with HRP (ref: A16096) was used for secondary staining at a dilution of 1:10,000 for 1 hour at room temperature. The membrane was washed three times in Tris-buffered Saline with 0.1% Tween after secondary antibody staining. Clarity Western ECL Substrate (ref: 1705060) was added to the blot and incubated at room temperature for 5 minutes. Western blots were visualized on a ProteinSimple FluorChem R imager.

### APEX2 proximity biotin labeling in stable HEK293T cell lines

The APEX2 proximity labeling methods we employed were adapted from Hung, *et al* [28]. All APEX2 proximity labeling was performed on the same day, in triplicate for each sample. HEK293T APEX2 stable cell lines were grown for 48-72 hours and supplemented with fresh media the morning of labeling. Biotin-labeling was initiated by changing the medium to fresh prewarmed medium containing 500 μM biotin-phenol. Plates were incubated for 30 minutes at 37 °C with 5% CO_2_. Cells, which detached during biotin-phenol incubation, were collected in a 50 mL conical tube, and 100 μL H_2_O_2_ was added to the media to achieve a final concentration of 1 mM to cells and incubated for 1 minute at room temperature. Cells were spun at 125 x g, and the supernatant was removed. Cells were washed twice with quencher solution (10 mM sodium ascorbate, 10 mM sodium azide, and 5 mM Trolox, in DPBS, freshly made) and twice with PBS. For each wash, 10 mL Quencher solution or PBS was added to cells, spun down at 125 x g, and removed the supernatant. The final pellets were flash frozen and stored at -80 °C.

### Streptavidin Pull-down of Labeled Proteins

Biotin-labeled cell pellets were thawed on ice and lysed with 500 μL of Lysis Buffer (150 mM NaCl, 1% Triton X-100, 0.5% Sodium Deoxycholate, 0.1% SDS, 50 mM Tris HCl pH 8.0, 10 mM ascorbate, 5 mM Trolox, Roche cOmplete Mini EDTA-free Protease Inhibitor Cocktail (ref: 11836170001)). Lysate was spun at 12,000 x g, and the supernatant was collected. 50 μL of Dynabeads MyOne Streptavidin T1 (ref: 65601) were equilibrated with Lysis Buffer for each replicate. Clarified lysate was added to equilibrated beads and nutated for 2 hours at 4 °C. A total of six bead washes were performed following removal of the unbound supernatant; extensive washing was started with Lysis Buffer, following which, RIPA (150 mM NaCl, 1% Triton, X-100 0.5% Sodium Deoxycholate, 0.1% SDS, 50 mM Tris HCl pH 8.0), KCl Buffer (1 M KCl, 50 mM Tris HCl, 5 mM EDTA, pH 8.0), 0.1 M Na2CO_3_, Urea Buffer (2 M Urea, 10 mM Tris HCl, pH 8.0), and repeating the last wash with RIPA. Following bead washes, 150 μL of Elution Buffer (50 mM Tris HCl, 2% SDS, 2 mM DTT, 10 mM EDTA) was added to the beads and samples were subsequently boiled at 95 °C for 15 minutes. The resulting supernatant was collected and processed for mass spectrometry analysis.

### Sample preparation for Mass Spectrometry

Following enrichment of labeled proteins using streptavidin beads, the eluate was processed using E3filters following the protocol reported recently [29]. In brief, the eluate samples were first boiled with final concentration of 10 mM Tris (2-carboxyethyl) phosphine and 40 mM chloroacetamide for 10 minutes at 95 °C. Then four volumes of Binding Solution (80% acetonitrile) were subsequently added, and this mixture was loaded onto the E3filters and centrifuged at 3,000 rpm for 2 min. The column was washed three times with Binding Solution, centrifuging and removing the wash solution each time. For digestion, Sequencing Grade Modified Trypsin (ref: V5111) was added at a 50:1 (wt/wt) ratio in 200 μL of Digestion Buffer (50 mM Triethylammonium bicarbonate) and incubated at 37 °C overnight with gentle shaking. Subsequently, the initial sample elution was collected, and the column was further washed with 200 μL Elution solution II (0.2% formic acid in water) and 200 μL Elution solution III (0.2% formic acid in 50% acetonitrile). The eluents were pooled and dried using a SpeedVac. Digested peptides were then desalted using C18 StageTips (CDS Analytical, Oxford, PA), and stored under -80 °C until further analysis.

### LC-MS/MS, protein identification, and bioinformatics analysis

Protein digests were analyzed using an Ultimate 3000 RSLCnano system coupled with an Orbitrap Eclipse mass spectrometer and field asymmetric ion mobility spectrometry (FAIMS) Pro Interface (Thermo Scientific) following procedures described previously [29]. The peptides were first resuspended in LC buffer A (0.1% formic acid in ACN), and half of the resuspension was analyzed using DIA method and the other half was analyzed using DDA method. For LCMS, the samples were first loaded onto a trap column (PepMap100 C18,300 μm × 2 mm, 5 μm particle; Thermo Scientific) and then separated on an analytical column (PepMap100 C18, 50 cm × 75 μm i.d., 3 μm; Thermo Scientific) at a flow of 250 nl/min. A linear LC gradient was applied from 1% to 25% mobile phase B (0.1% formic acid in ACN) over 125 min, followed by an increase to 32% mobile phase B over 10 min. The column was washed with 80% mobile phase B for 5 min, followed by equilibration with mobile phase A for 15 min.

For data-dependent acquisition (DDA) MS analysis, the detector type was Orbitrap with resolution of 60,000; the MS1 scan range was set to 375-1500 m/z, AGC target was set to 4E5, and the maximum injection time mode was set to 50 ms. For MS2 analysis, precursors with charge states 2–5 were selected. The isolation mode was Quadrupole, collision was by HCD at 30% normalized collision energy (NCE). The Orbitrap was set to detect MS2 fragments at 15,000 resolution; the AGC target was set to 5E4, and max injection time 22 ms. Dynamic exclusion was set to 30 s. Monoisotopic precursor selection (MIPS) was set to Peptide. For FAIMS settings, a 3-CV experiment (−40|-55|-75) was applied.

For data-independent acquisition (DIA) MS analysis, the detector type was Orbitrap with resolution of 120,000; the MS1 scan range was set to 380-985 m/z, AGC target was set to 4E5, and the maximum injection time mode was set to Auto. The isolation mode was quadruple; DIA window type was auto, and isolation window (*m/z*) was 8 with an overlap of 1. The detector type for MS/MS was Orbitrap with a resolution of 15,000; normalized automatic gain control target (%) was 800, and the maximum injection time mode was auto; and the loop control was 2 s. The FAIMS settings were the same as DDA mode.

The DDA raw data were processed using MaxQuant (version 2.4.9.0), and the DIA raw data were processed using Spectronaut software (version 19.0), following similar procedures described previously [29,30]. In MaxQuant Group-specific parameters, the Type was set to FAIMS, and the LFQ was enabled with minimum ratio count of 1. The other parameters were taken default settings, similar to the procedures described previously [29]. In Spectronaut, a library-free DIA analysis workflow with directDIA+ was applied. Detailed parameters for Pulsar and library generation were similar to the settings reported recently [30]. Bioinformatics analyses including *t* test, correlation, volcano plot, and clustering analyses were performed using Perseus software (version 2.0.11) and GraphPad Prism (version 11) unless otherwise indicated.

## Results and Discussion

### Quantitative comparison of DIA and DDA proximity proteomics

Protein complexes derived from proximity labeling pulldown experiments have been commonly analyzed using DDA MS due to its simplicity in data interpretation and the established bioinformatics pipelines [31]. On the other hand, DIA MS is rapidly gaining popularity due to its higher reproducibility, superior quantitative accuracy, and deeper coverage compared to traditional DDA MS [32,33]. However, to the best of our knowledge, direct comparisons of the two acquisition techniques in the context of proximity proteomics have not been reported before, although such comparative studies have been carried out using other experimental systems, such as cross-linking MS [34] and whole cell proteomics [35]. Herein, we fused APEX2 to either the N- or C-terminus of the TRMT1 bait and acquired MS data using both DIA and DDA methods on the same set of proximity labeling samples, in order to perform head-to-head comparisons. An illustration of the experimental workflow is shown in **Figure 1**.

**Figure 1.**
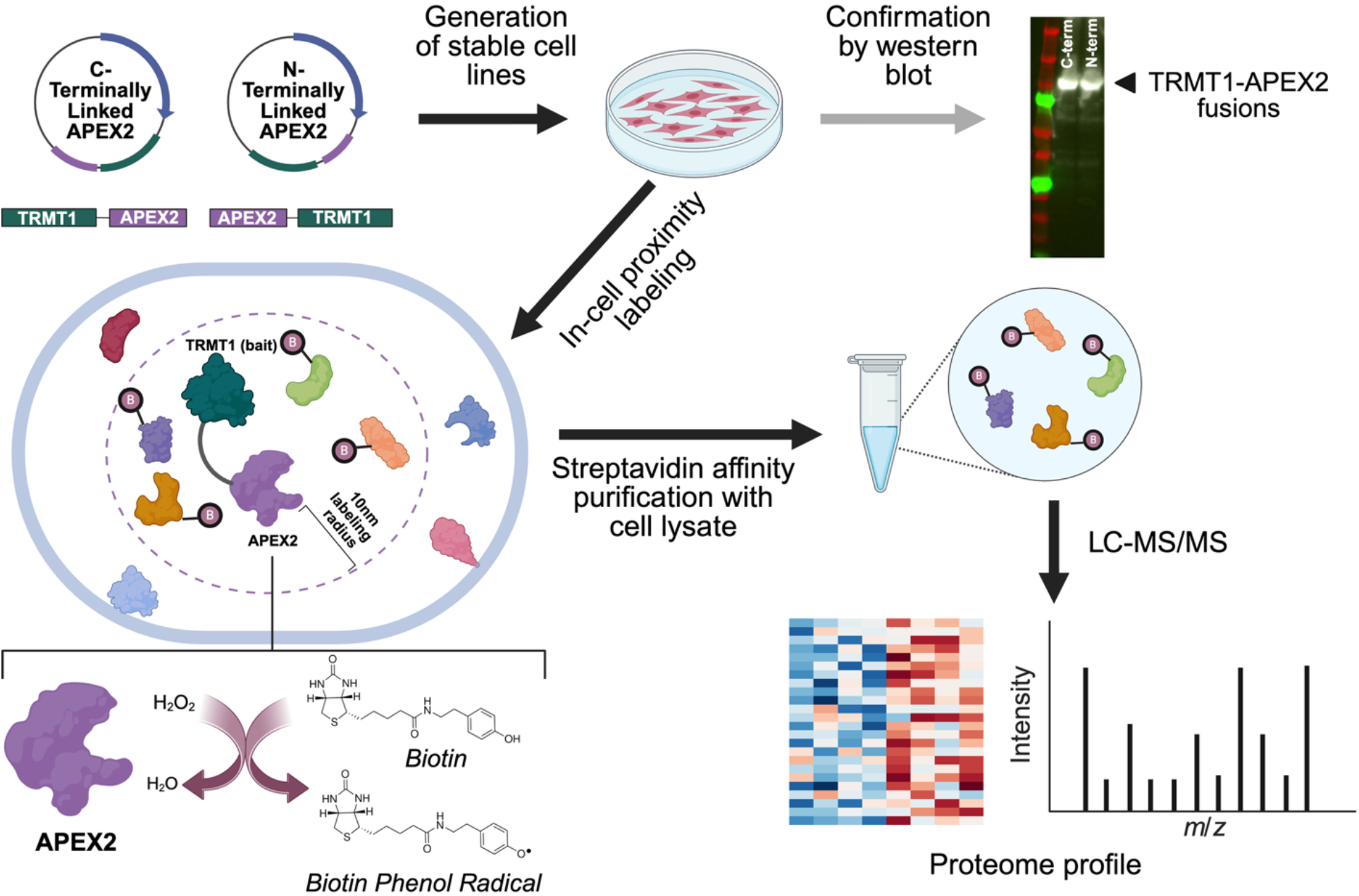
Illustration of the dual proximity labeling workflow to study the TRMT1 interactome. N- or C-terminally APEX2 fusions with TRMT1 are generated, transfected to make stable cell lines, and TRMT-APEX2 expression confirmed by Western blot. In-cell proximity labeling tags nearby proteins with biotin, which are pulled down and analyzed by LS-MS/MS to identify and map the TRMT1 proximal proteome. Image created with BioRender.com.

Our data indicated that DIA MS can identify approximately 8,000 proteins from each of the three pulldown complexes (i.e., N-terminal APEX2-TRMT1 fusion, C-terminal TRMT1-APEX2 fusion, and APEX2 alone) (**Figure 2A**). The three biological replicates of each pulldown complex showed generally good reproducibility, averaging 15-20% coefficient of variation (CV) and Pearson correlation 0.95 or higher, except for the N-term fusion pulldown where a possible outlier slightly skewed the performance (**Figure 2, B-C**). Qualitatively, the three replicate experiments also showed significant overlaps, sharing 90% or more commonly identified proteins (**Figure 2D**). By contrast, the DDA method reported on average 3,900 protein identifications from the respective pulldown experiments, two times less than the data from the DIA method. DDA also showed larger variabilities among the replicate experiments (**Figure 2, G-J**). Previously, Khalil *et al*. reported nearly 45% more hits when comparing DIA with DDA method in the context host cell protein analysis [36]. We also previously noticed approximately 40% more proteins when analyzing the yeast whole cell proteome using DIA and DDA method [37]. Along these lines, the findings from this study show that DIA MS can remarkably facilitate proximity proteomics analysis and offer more reproducible and much deeper coverage in comparison to DDA method.

**Figure 2.**
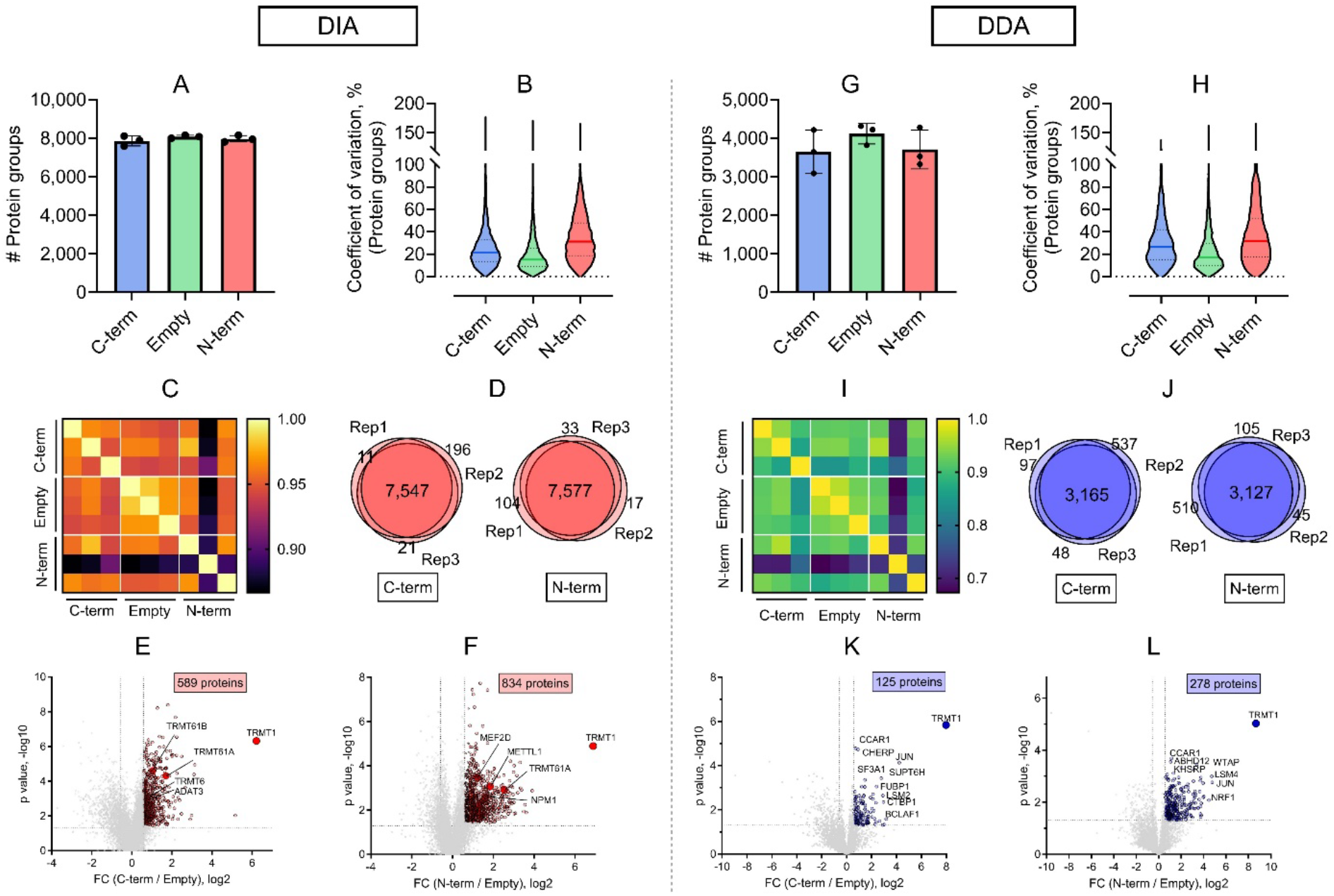
Quantitative comparison of DIA and DDA proximity proteomics. (A) Number of protein identifications derived from the DIA analysis of the C-terminal, empty vector (genitive control), and N-terminal pulldown experiments. (B) Protein group coefficient of variation analysis. Bold lines indicate the median values. (C) Pearson correlation analysis among the replicates of the three pulldown experiments. (D) Overlapping analysis of the three replicates among the C-terminal and N-terminal pulldown experiments. (E-F) Volcano plot of the proximity labeling enrichment proteins. Fold change (FC) cutoff is 1.5 and the adjusted p value is 0.05. (G-L) Similar to DIA analysis, the plots show the DDA data analysis results.

To determine the potential interacting proteins to the bait protein TRMT1, we performed two-sample t-test of the C-or N-terminal fusion pulldown complex with a negative control, where HEK293 cells expressing APEX2 alone were treated with biotin. We required potential interaction candidates to meet fold change (FC, > 1.5) and adjusted p value (< 0.05) cutoffs, while also being matched with at least two unique peptides. Based on these criteria, 125 and 278 proteins derived from DDA data were identified as significantly enriched hits from the C- and N-terminal fusion complexes, respectively (**Figure 2, K-L**), whereas three to four times more significant proteins were identified by DIA method (**Figure 2, E-F**). The above data indicated again that DIA MS not only can provide a deeper proximity proteome, but also can detect more potential interacting proteins than conventional DDA MS methods.

### Assessment of N-terminal and C-terminal proximity proteome

Because proximity labeling enzymes have a finite labeling radius (for instance, < 20 nm for APEX2), fusing to different termini (N- or C-) of a bait protein can position APEX2 differently within a complex or organelle, resulting in unique snapshots of the local environment with specific, and sometimes distinct, interaction networks [38,39]. Therefore, when studying a new protein, it is often recommended to generate both N-terminal and C-terminal fusion constructs to maximize the coverage of the interactome and ensure that protein localization or function is not perturbed by the tag [40]. In this study, we fused APEX2 to either N- or C-terminus of TRMT1 protein, and carried out the pulldown and LCMS analyses in parallel along with a negative control experiment. Such a comprehensive head-to-head comparison dataset is useful in determining the proximal and interacting proteins of the bait TRMT1. Our data suggest that the N-terminal and C-terminal fused TRMT1 constructs capture largely similar proximity proteomes. Nearly 94% or higher of the identified proteins were shared by both N- and C-terminal APEX2 pulldowns, and their quantities were also generally equivalent (**Figure 3, A-B**). This finding indicates that the two termini of TRMT1 protein may be surrounded with similar protein complexes and that APEX2 labeling does not significantly perturb protein interactions of localization in a termini-specific way.

**Figure 3.**
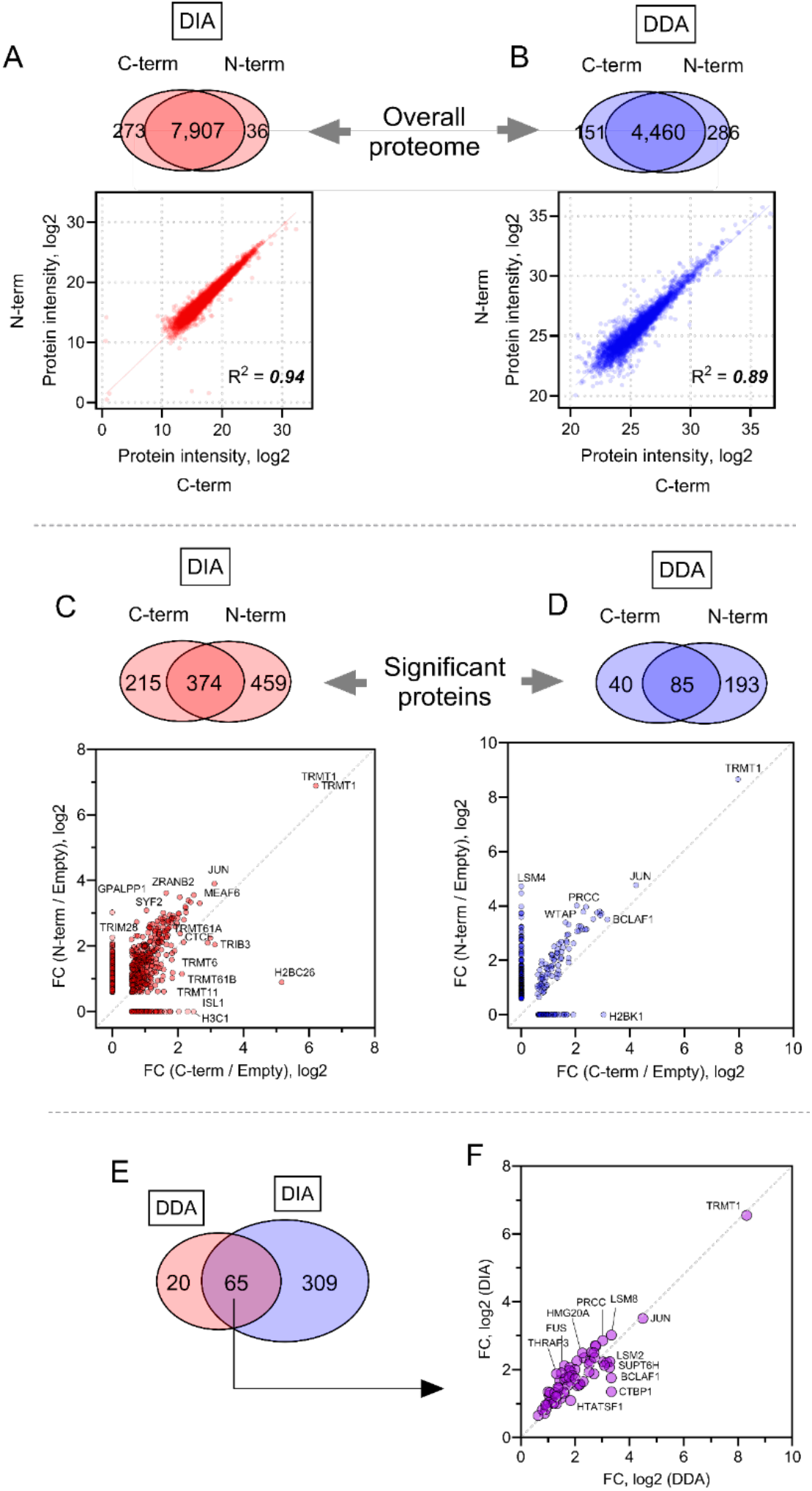
Assessment of N-terminal and C-terminal proximity proteome. (A-B) Venn diagrams show the overlapping analyses of the overall proteomes obtained from the two terminal pulldown experiment using DIA and DDA method, respectively. The two correlation plots show the correlation of the median value of the three replicates in each pulldown experiment. (C-D) Venn diagrams show the overlapping analyses of the proteins that are significantly enriched in the two terminal pulldown experiment compared to the negative control using DIA and DDA method, respectively. The two correlation plots show the FC correlation of the overlapped proteins (374 and 85 proteins, respectively). (E) Overlapping analysis of the total significant proteins derived from the DIA and DDA method, respectively. (F) FC correlation of the overlapped proteins (65 proteins in total) between the two methods.

From the DIA data, we found that 834 and 589 of the total 7,815 proteins identified with at least two unique peptides were significantly enriched (fold change > 1.5; adjusted *p* < 0.05) from the N-terminal and C-terminal pulldown complexes, respectively, when compared with the APEX2-only control (**Figure 2, E-F**). Among these significant protein hits, 374 of them showed the same enrichment pattern from both N- and C-terminal pulldown complexes, whereas 459 and 215 proteins showed distinct trends in the N-versus C-terminal APEX2 pulldowns, respectively (**Figure 3C**).

Gene Ontology (GO) enrichment analysis (**Figure 4, A-D**) of the overall 1,048 significantly enriched proteins in the DIA data set revealed that a majority of them were localized in the nucleoplasm (58%) or nucleus (72%). TRMT1 has both nuclear and mitochondrial localization signals and subcellular fractionation experiments have shown that it is highly enriched in the nucleus and mitochondria, with some presence in the cytoplasm as well [22]. TRMT1 may also undergo re-localization from mitochondria and the cytoplasm to the nucleus upon neuronal activation [41]. This is broadly consistent with the significant nuclear localization of the 1,048 candidate proximity proteins from our data. Furthermore, the biological process (BP) enrichment analysis of these 1,048 protein hits showed extremely high representation of proteins involved in mRNA processing (*p* = 4.11 × 10^−85^) and RNA splicing (*p* = 3.42 × 10^−70^). GO enrichment analysis of the DDA data revealed similar patterns regarding cellular compartment and biological process (BP), although the numbers of significant proteins from the two terminal pulldown experiments were low (**Figure 2, K-L; Figure 3D**). Together, the strong enrichment of TRMT1-proximal proteins identified in the nucleus and with involvement in RNA processing and splicing suggest that TRMT1 may play additional roles outside of canonical tRNA modification and could be involved in complexes that mediate or coordinate nuclear RNA processing events.

**Figure 4.**
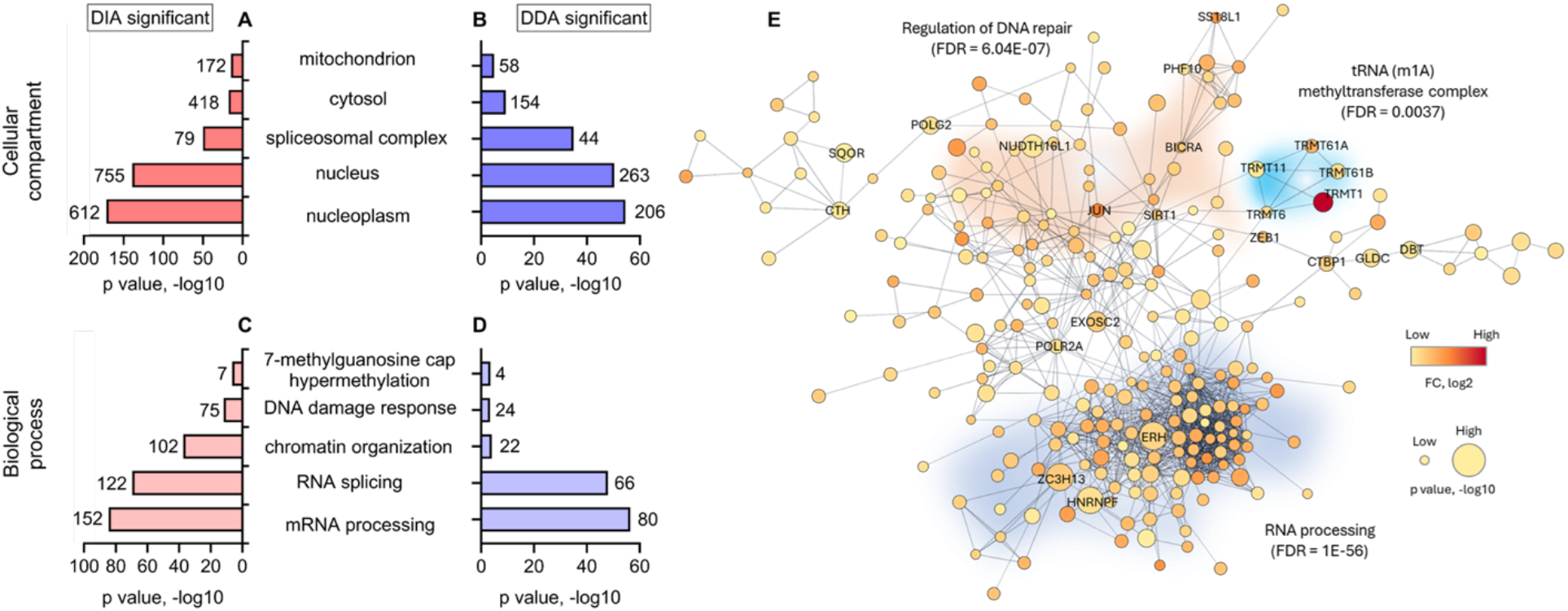
Dissection of TRMT1 interactome. **(A-B)** Gene Ontology cellular compartment enrichment analysis of the significantly enriched proteins from the DIA and DDA method, respectively. The top five over-represented terms were plotted here. The numbers on top of each bar indicate the number of proteins in the respective terms. **(C-D)** Similarly, the biological process terms were plotted. **(E)** Predicted protein interaction network by the STRING database. The total 394 significantly enriched proteins by the two acquisition methods (as shown in Figure 3E) were imported into Cytoscape (version 3.10.4) using a confidence score cutoff of 0.7, with the ‘load enrichment data’ option enabled. Singleton nodes were excluded from the network. A few representative protein clusters with their FDR values are highlighted.

### Dissection of TRMT1 interactome

Although systematic profiling of TRMT1-associated proteins has been limited and the GO analysis above was unexpectedly enriched in nuclear RNA processing proteins, several significantly enriched candidates in our DIA dataset are biologically consistent with the known role of TRMT1 in tRNA modification and translation (**Figure 2, E-F**). Notably, we identified multiple proteins involved in tRNA modification, including TRMT6, TRMT61A, TRMT61B, METTL1, ADAT3, DUS1L, PUS7L, and TRMT11 [24,42–47]. The TRMT6/61A complex catalyzes tRNA m1A58 formation and METTL1 is the catalytic core of the tRNA m7G methyltransferase complex [24,42]. Likewise, ADAT3, DUS1L, and PUS7L each carry out distinct tRNA editing or modification reactions [44–46]. None of these proteins are established TRMT1 binding partners, but their enrichment in our dataset supports the localization of TRMT1 near other tRNA modification enzymes. Furthermore, we detected enrichment in some RNA polymerase III-associated proteins, including POLR3F, POLR3G/POLR3GL, and BDP1, which suggests proximity between TRMT1 and proteins or pathways involved in tRNA transcription or tRNA biogenesis [48–50]. Together, these candidate hits provide biologically plausible examples of TRMT1-proximal proteins that align with established roles for TRMT1 in tRNA methylation and translation control. Notably, the DDA MS dataset either did not detect or show significance for the majority of these candidate proteins, suggesting that the lack of proteome depth is detrimental to proximity labeling-based interactome analysis.

In addition to the tRNA modification and RNA metabolism factors above, enriched TRMT1-proximal candidates also included proteins linked to redox regulation and cellular stress response, including SLC7A11, GPX1, PRDX4, PRDX6, ACO1, CBS, CTH, MPST, and SQOR [51–61]. These redox-associated candidate hits are notable because TRMT1-catalyzed tRNA modifications are required for redox homeostasis and oxidative stress response. Their enrichment here suggests possible further functional connections between TRMT1 and adaptation to cellular redox stress.

STRING network analysis of the significantly enriched DIA candidates revealed several interconnected protein networks that were consistent with the GO enrichment results (**Figure 4E**). Prominent clusters include tRNA methylation, RNA processing, and DNA repair, reflecting nuclear environments where TRMT1 is localized. Similarly, comparing our DIA MS candidate list with the BioGRID database for protein interactions, we find reported TRMT1 associations for significantly enriched protein hits including ACADVL (involved in mitochondrial redox/metabolism), TRMT61A, EFTUD2 and SART3 (involved in RNA processing), and RPA1-3 (involved in DNA replication and repair) [42,62–66].

## Conclusion

Integrating APEX2 proximity labeling with advanced DIA MS acquisition techniques enabled sensitive mapping of the TRMT1-proximal interactome in human cells. By comparing APEX2 fusions at both termini of TRMT1 (dual-labeling), we captured a broad and largely overlapping set of candidate proximity proteins. Direct comparison of DIA and DDA acquisition showed that DIA provided much greater depth and sensitivity, and this method likely allows for the detection of transient and weak protein interactions that may be missed by traditional DDA methods. The dataset of enriched TRMT1-proximal candidates includes many nuclear RNA processing factors and biologically plausible protein associations connected to tRNA modification, tRNA biogenesis, and redox / cell stress pathways. Although further validation will be required to determine which proteins represent direct or functional TRMT1 interactors, this study provides a resource for future investigation of TRMT1 regulation and function, and provides a comprehensive head-to-head comparison of DDA and DIA MS methods in proximity-based proteomics.

## Acknowledgments

This work was supported by the US National Institutes of Health, National Institute of General Medical Sciences, under awards R35 GM143000 to JSM, T32 GM133395 CBI fellowship to AD and HB, and P20 GM104316 that funded Orbitrap Eclipse mass spectrometry and other key instrumentation used in this study. The content is solely the responsibility of the authors and does not necessarily represent the official views of the National Institutes of Health.

## Notes

### Competing Interest Statement

The authors have declared no competing interest.

